# A negative binomial latent factor model for paired microbiome sequencing data

**DOI:** 10.1101/2024.12.01.626246

**Authors:** Hyotae Kim, Nazema Siddiqui, Lisa Karstens, Li Ma

## Abstract

**Motivation:** Microbiome compositional data are often collected from several body sites and exhibit dependency among them. Analyzing microbial compositions from different sites jointly allows for effective borrowing of information by exploiting the underlying cross-site correlation, which can lead to more effective statistical analysis, especially when the sample size at one or both sites is limited. To this end, we introduce a joint model for microbiome compositions at two (or more) sites within the same subjects. Our model incorporates (i) latent factors shared across two body sites to explain the common subject effects and to serve as the source of correlation between the two sites; and (ii) mixtures of latent factors to allow heterogeneity among the samples in their level of cross-site association. The model is illustrated with synthetic data and we apply it in a case study involving samples of the urinary and vaginal microbiome collected from women.

**Results:** Simulation studies show how common subject effects influence regression analysis results; a stronger association between two sites in the data causes a greater degree of bias in the analysis. The model with latent factors mitigates the bias present in the model without latent factors, whereas the two models perform comparably for the data set without paired associations. In a case study involving samples collected from a study on the female urogenital microbiome with aging (e.g., the UMICRO study), our model leads to the detection of covariate associations of the vaginal and urinary microbiome composition that are otherwise not statistically significant under a similar regression model applied to the two sites separately. Our model also enables prediction of the microbial abundance at one site based on observations from another site. We also consider a model extension that allows the clustering of subjects (samples) and cluster-specific levels of paired association. Under the extended modeling framework, the clusters can be classified according to their association strengths.

## 1 Introduction

The advent of next-generation sequencing (NGS) technology enables the identification of a wide range of microbes from environmental samples without the need for cultivation, thus facilitating the exploration of microbial communities. The two most widely used methods for sequencing microbial communities are universal marker gene amplicon sequencing, such as the 16S rRNA gene, and whole metagenome shotgun sequencing (WMS) of all microbial genomes. These methods produce sequencing reads, which are subsequently mapped to taxa at various taxonomic levels using bioinformatic preprocessing pipelines, such as DADA2 [Callahan et al., 2016]and MetaPhlAn [Blanco-Míguez et al., 2023]. As a result of this taxonomic profiling, a large, highly sparse count table of taxa per sample is produced, with typically a finer taxonomic resolution in WMS than in 16S amplicon sequencing.

To gain a comprehensive understanding of the human microbiome, researchers often collect data from multiple body sites for each individual and compare the composition and functions of the microbial communities in different parts of the body (e.g., [HMPC, 2012a, HMPC, 2012b]). The UMICRO data set, which we will use as a case study, includes vaginal-urine paired samples obtained by vaginal swabbing of the distal vagina and urine collection by transurethral catheterization, with the aim of identifying the microbial composition of the two communities and analyzing their variations with respect to the menopausal status of the participants. As the sample pairs were collected from two different body sites of the same subject, common effects associated with the subject are likely to be present in both vaginal and urine samples. This data set motivates the development of a model to capture the potential associations between the vaginal and urine niches.

For the analysis of microbial compositional data in the form of a count table with rows for samples and columns for taxa, modeling methods originally developed for RNA sequencing (RNA-seq) or single-cell RNA sequencing (scRNA-seq) data can be adopted. Similarly to microbial compositional data, RNA sequencing data provide sparse count tables that report the number of sequence fragments assigned to each gene per sample or cell, for which the following models have been devised: negative binomial models [Robinson et al., 2010, Love et al., 2014],zero-inflated negative binomial models [Risso et al., 2018], Poisson zero-inflated log-normal models [Wang et al., 2018],and truncated Gaussian hurdle model [Finak et al., 2015]. In addition, [Martin et al., 2020, Morton et al., 2019, Paulson et al., 20 Sohn et al., 2015]introduced beta-binomial, multinomial, and zero-inflated Gaussian models, which were specifically developed for microbiome data. Although zero-inflated models were created to accommodate the pronounced sparsity of microbiome data, their validity in count-based models for sequencing reads remains controversial. As discussed in [Silverman et al., 2020, Sarkar and Stephens, 2021], in many common settings involving sparse sequencing count data, the abundance of zeros can often be adequately accommodated by simply incorporating overdispersion into count-based sampling models, such as in negative binomial models, without needing an additional zero-inflation component. We share this viewpoint, and because we are considering overdispersed count-based models in this paper, we do not by default incorporate an additional zero-inflation component, but in cases where such a component is indeed justified, incorporating it into the model is straightforward.

We propose a model-based approach to jointly account for microbiome compositions at two (or more) sites, adopting a negative binomial regression model for each site while incorporating a shared latent factor to parsimoniously capture potential correlation in paired samples from different body sites in some taxa. Specifically, two negative binomial distributions, one for each site, share a set of latent factors, which are interpreted as unobserved common effects that contribute to their correlation. Under suitable choices of priors on the model parameters, posterior samples can be drawn efficiently using a Gibbs sampler that employs a data-augmentation technique called Pólya-Gamma augmentation. We extend our model on the latent factors to accommodate the common assumption that the samples are not homogeneous but rather form subgroups or clusters, each with their own cross-site correlation patterns. We present versions of the model that work both when the subgrouping is observed and when it is not. In the latter case the subgrouping is inferred from the data. We carry out a case study on the UMICRO data in which the study subjects come from three subgroups by study design.

The rest of the paper is organized as follows. Section 2 introduces our negative binomial latent variable model with Section 2.1 for the base modeling framework and Section 2.2 for some model extensions. In Section 2.3, we present a hierarchical representation of our model with prior specifications for the model parameters, followed by a posterior prediction approach. The proposed model is illustrated through synthetic data in Section 3.1 and the UMICRO study in Section 3.2. Finally, Section 4 concludes.

## 2 Methods

### 2.1 Joint Negative Binomial Model

For a given taxon (e.g., genus), we use *y*_*si*_ to denote the observed count in sample *i* at body site *s*, with *i* = 1, ⋯, *n* and *s* = 1, 2. Note that the data are actually also indexed by taxa, but for simplicity, throughout our description of the model and the computational recipes, we suppress the index for taxa, as our model is taxon-specific and is applied to each of the taxa separately. Let NB(*μ, α*) be a negative binomial distribution with mean *μ* and variance (1 + *μα*)*μ*. We consider the following joint negative binomial model (JNBM) that relates the counts from two sampling sites through a taxon-specific latent factor, *γ*_*i*_, in the form of a multiplicative factor on the mean count for the taxon,

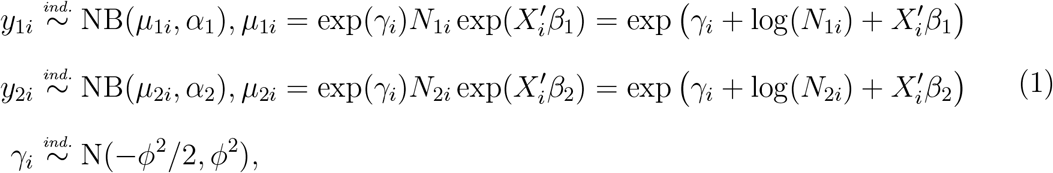

where *X*_*i*_ denotes the vector of covariates for sample *i*, and *β*_*s*_ is the regression coefficient vector for body site *s. α*_*s*_ is the overdispersion parameter of the negative binomial distribution. *N*_*si*_ indicates the total number of read counts, that is, the sum of *y*_*si*_ over all taxa of interest. The latent factor *γ*_*i*_ (or its exponential exp(*γ*_*i*_))is an unobserved variable that represents a site-invariant sample-specific effect. The multiplicative effect of exp(*γ*_*i*_) on the mean, *μ*_*si*_, has E(exp(*γ*_*i*_)) = 1 and Var(exp(*γ*_*i*_)) = exp(*ϕ*^2^) *−* 1. This distributional assumption with the mean of 1 enables the random effects of exp(*γ*_*i*_) to have no preference for a positive impact (*>* 1) or negative impact (*<* 1). The parameter *ϕ*^2^ measures the strength of the association between the two body sites; when *ϕ*^2^ becomes zero, the model is reduced to two independent negative binomial distributions for each body site, that is, NB for *s* = 1, 2. We will refer to it as the separate negative binomial model (SNBM). Section 3.2 will compare our joint two-site model with SNBM in a case study on the UMICRO data.

To perform JNBM-based Bayesian inference, we need priors for (*α*_*s*_, *β*_*s*_, *ϕ*^2^); a description of the complete Bayesian hierarchical model can be found in Section 2.3.1. A fully conjugate sampling recipe for posterior inference based on the Pólya-Gamma data augmentation technique is detailed in Supplementary Material B.

For brevity in the model description, we assume balanced paired data, i.e., both sites have the same number of samples. However, the model can also be used for unbalanced data where some samples are available only for one site.

### 2.2 Model Extension

Although JNBM assumes a constant level of cross-site association throughout all paired samples, in real-world examples, this assumption is often unrealistic. However, allowing each sample to have its own level of association may lead to an overly flexible model. We thus compromise and assume that there are subgroups (either observed or unobserved) among the samples for which the extent of cross-site association is comparable. Consequently, we extend the base joint model by allowing different sample groups to exhibit different levels of paired association, which can easily be achieved by replacing the normal distribution for the latent factors with a mixture distribution. The following is a two-component normal mixture-based model, which randomly assigns paired samples to two groups:

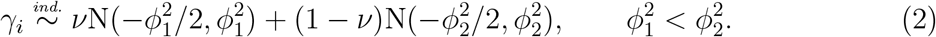

It should be noted that we enforce an inequality constraint on the mixing parameter, 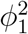 and 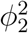, to ensure model identifiability. The general form of the mixture prior is given by 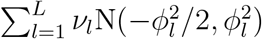 with 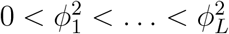 and 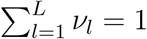 for *L >* 1. In the paper, we will call this joint negative binomial model with the two-component mixture distribution JNBM Mix. Supplementary Material A.4 presents JNBM Mix’s superior predictive performance over JNBM, along with random clustering results for Lactobacillus samples as an example.

The clustering of paired samples can also be accomplished by using a given variable *g*(*i*) = *{*1, ⋯, *L}* rather than random assignment with *{ν*_*l*_*}*, where differences in association levels across *L* groups may serve as an indicator of the proximity among them. The mixture distribution for *γ*_*i*_ is adjusted as follows:

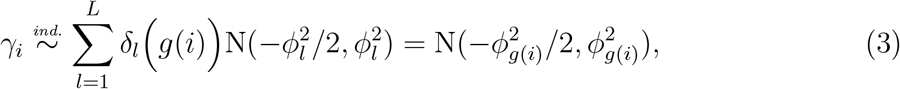

where *δ*_*l*_(*x*) is a delta function with *δ*_*l*_(*x*) = 1 for *x* = *l* and 0 otherwise. For the UMICRO case study, we use Study Group variable for clustering, such that *g*(*i*) = *{*Postmenopausal with no estrogen, Postmenopausal on estrogen, Premenopausal*}*, which model is named JNBM SG. Figure S11 shows the posterior distributions of 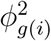 for each study group under the model, revealing obvious similarities between the two postmenopausal groups and differences in location or/and dispersion between postmenopausal and premenopausal for *Lactobacillus, Aerococcus, Prevotella* and *Corynebacterium*. JNBM SG also outperforms SNBM and JNBM in prediction, which will be discussed in Section 3.2 and Supplementary Material A.2.

### 2.3 Posterior inference

#### 2.3.1 Hierarchical model representation

Below is a full description of the joint negative binomial model (JNBM) presented in (1) of Section 2.1. Let ***y*** = *{y*_*si*_ : *s* = 1, 2 and *i* = 1, ⋯, *n}*, ***γ*** = *{γ*_*i*_ : *i* = 1, ⋯, *n}*, and *β*_*s*_ = *{β*_*sp*_ : *p* = 1, ⋯, *P}*. The model is defined as

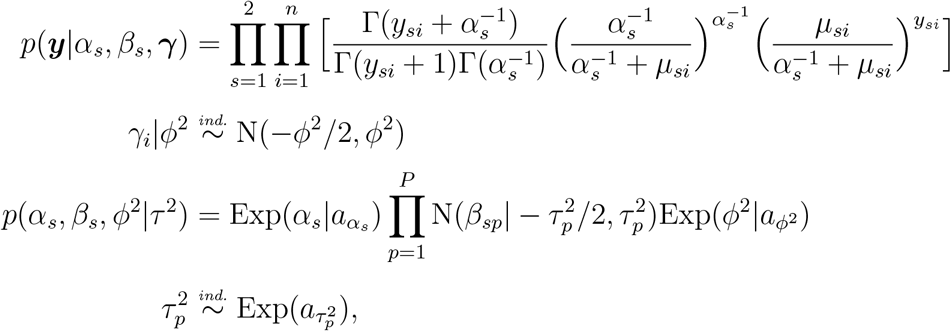

where 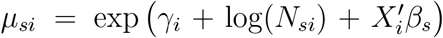. For simplicity, we place exponential priors on 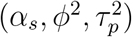 their rate parameters were chosen empirically, which are conservative choices and lead to substantial prior-to-posterior learning. For example, in Section 3.1, we set 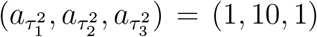 for a simulation study with true regression coefficients of (*β*_11_, *β*_12_, *β*_13_) = (0.05, 0.001, 0.03) and (*β*_21_, *β*_22_, *β*_23_) = (0.03, 0.002, 0.01). With the prior means of 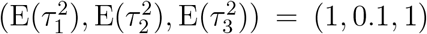, the 95% prior uncertainty bands for *β*_*s*1_, *β*_*s*2_, and *β*_*s*3_ are given by (*−*2.5, 1.5), (*−*0.7, 0.6), and (*−*2.5, 1.5), respectively. The conservative choices result in much tighter 95% posterior uncertainty bands with substantial shifts of the posterior means (from the prior means) toward the true values. Likewise, we chose conservative hyperparameters of 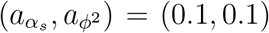. For the case study in Section 3.2, we set the hyperparameters to 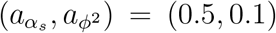, as well as 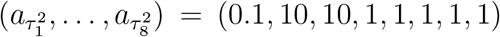 for coefficients of the intercept, two continuous covariates, and five binary covariates. We observed that this prior specification gives rise to such prior-to-posterior learning, although the true underlying parameter values are unknown in the case study. As in *γ*_*i*_, the regression coefficients *β*_*sp*_ are assigned normal distribution priors with mean 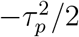 and variance 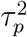, allowing the multiplicative effects of exp(*β*_*sp*_) on *μ*_*si*_ to have their means equal to 1 with their variances of exp 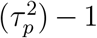. The following section discusses the augmented likelihood under the negative binomial modeling framework, which enables the regression coefficients and the latent factors to have full conditionals in closed form.

Similarly, we can represent the extended models – JNBM Mix and JNBM SG – with variations in distributional assumptions on *γ*_*i*_ as follows,

[**JNBM Mix**] : with auxiliary variables *{ξ*_*i*_*}* for a hierarchical representation of the mixture distribution on *γ*_*i*_,

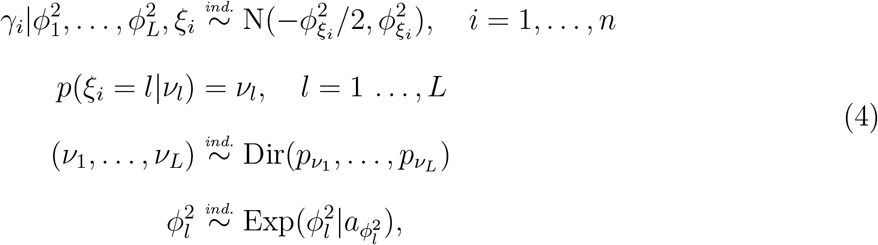

where Dir(*p*_1_, ⋯, *p*_*L*_) denotes a Dirichlet distribution with *p*_*l*_ = 1*/L*. [**JNBM SG**] :

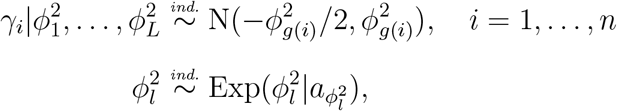

where *g*(*i*) is an observed variable, e.g., the study group clinical variable of the UMICRO data in Section 3.2. The hyperparameter 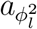 of the exponential prior for 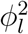 is chosen in the same manner as *a*_*ϕ*_^2^ of *ϕ*^2^ in JNBM.

#### 2.3.2 Cross-site prediction

In making predictive inferences about unknown observables *y*^*∗*^, we consider the posterior predictive distribution below, from which we draw predictive samples and compute their average. The posterior predictive distribution is

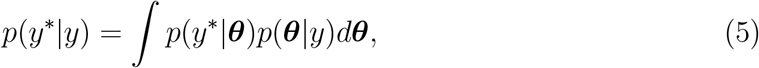

where *y* denotes the observed data and ***θ*** is a vector of model parameters: (*α, β*) for SNBM and (*α, β, γ*) for JNBM. Posterior predictive samples for *y*^*∗*^ are drawn from the negative binomial sampling distribution *p*(*y*^*∗*^|***θ***) with parameters that are substituted with the posterior samples of ***θ*** taken from *p*(***θ***|*y*). Then, the mean of the posterior predictive samples becomes the predictive value for *y*^*∗*^.

One useful feature of our joint model is the ability to predict counts at one site using observations from the other site. Suppose, for example, that *y*_1*i*_*′* for *i*^*′*^ *∈ I ⊂ A ≡ {*1, ⋯, *n}* are unobserved and prediction targets. In SNBM, posterior sampling of model parameters (*α*_1_, *β*_1_) is based only on observations *{y*_1*i*_ : *i ∈ A − I}*, while, in JNBM, not only *{y*_1*i*_ : *i ∈ A − I}* but also *y*_2*i*_*′, i*^*′*^ *∈ I*, are used for (*α*_1_, *β*_1_, *γ*_*i*_*′*) posterior sampling, specifically for latent factors *γ*_*i*_*′*. By plugging the posterior samples of the model parameters into the sampling distribution NB 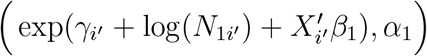, with *γ*_*i*_*′* = 0 for SNBM, we can draw posterior predictive samples for *y*_1*i*_*′*. Let 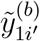 be a posterior predictive sample from NB 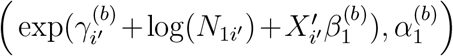 with 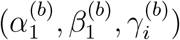 acquired at *b*-th MCMC iteration for *b* = 1, ⋯, *B*. Then, the predictive value 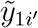 for *y*_1*i*_*′* is defined as the posterior mean, that is, 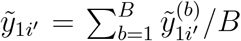. Section 3.2 discusses the predictive performance of the models, which is based on the leave-one-out cross-validation (LOOCV) approach; we select one observation from the 2*n* observations across *n* pairs of samples as a test set (i.e., an unknown observable for prediction), fit the models to the remaining data, and predict the test set value. We repeated the LOOCV 2*n* times to obtain the predictive values for all sample pairs of a given taxon.

### 3 Results

### 3.1 Simulation

We simulate paired samples with varying levels of association, comparing separate and joint negative binomial models to empirically grasp the role of latent factors in the joint model. Synthetic data were generated by drawing *n* = 300 pairs of samples from negative binomial distributions with latent factors defined as,

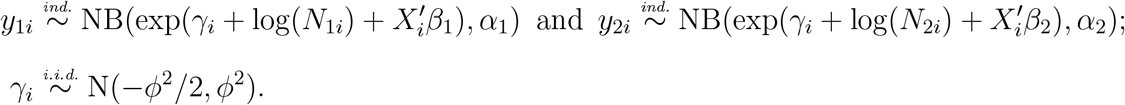

The dispersion parameters and regression coefficients are arbitrarily chosen to be *α*_1_ = 2, *α*_2_ = 1, *β*_1_ = (*β*_11_, *β*_12_, *β*_13_) = (0.05, 0.001, 0.03), and *β*_2_ = (*β*_21_, *β*_22_, *β*_23_) = (0.03, 0.002, 0.01),= where *X*_*i*_ = (*x*_*i*1_, *x*_*i*2_, *x*_*i*3_)^*′*^ is a vector consisting of an intercept *x*_*i*1_, a continuous covariate *x*_*i*2_ *∼* Unif(20, 80) and a binary covariate *x*_*i*3_ *∼* Bern(0.3). Unif(*a, b*) and Bern(*p*) are (continuous) uniform and Bernoulli distributions with means (*a* + *b*)*/*2 and *p*, respectively. Again, the dispersion parameter *ϕ*^2^ of the latent factors in the underlying distribution determines how strong the association between pairs of samples is, and it is set to 0 for independent pairs and to 2 or 10 for dependent pairs, with 10 indicating a stronger association. *N*_1*i*_ and *N*_2*i*_ are offsets, which in real-world applications are used to normalize for sequencing depth (or the sum of abundances across selected taxa of interest). We set 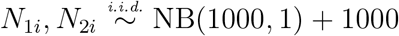 sampling of *N*_*si*_ from the negative binomial distribution results in count data with a right-skewness as with the sequencing depths in the UMICRO data set (but on a smaller scale). By repeatedly sampling with the parameters, covariates, and offsets defined above, we produced 300 sets of 300 sample pairs for the following analysis.

We evaluate the separate and joint models with regard to the accuracy of their estimated regression coefficients. Table 1 shows the mean square error (MSE) between the estimated and true coefficients for the two covariates under each model, demonstrating that JNBM is superior to SNBM for the data with non-zero associations (*ϕ*^2^*≠* 0). This indicates that ignoring the latent sample effect (*γ*_*i*_) as in SNBM can severely reduce the efficiency in estimating the underlying relationships between the response variables and predictors. As expected, the difference in estimation accuracy between the two models becomes greater as the paired association in the underlying distribution becomes stronger, and therefore the common effects become larger.

**Table 1.** MSE, denoted as 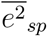 with site *s* = 1, 2 and covariate *p* = 2, 3 for regression coefficients *{β*_*sp*_*}*, with its standard error 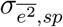 in parentheses.

| Scenario | Model | $\overline{e^2}_{12}(\sigma_{\overline{e^2},12}) \times 10^5$ | $\overline{e^2}_{13}(\sigma_{\overline{e^2},13}) \times 10^2$ | $\overline{e^2}_{22}(\sigma_{\overline{e^2},22}) \times 10^5$ | $\overline{e^2}_{23}(\sigma_{\overline{e^2},23}) \times 10^2$ |
| --- | --- | --- | --- | --- | --- |
| $\phi^2 = 0$ | SNBM | **1.53** (0.14) | 2.10 (0.18) | 0.80 (0.07) | **1.10** (0.09) |
|  | JNBM | 1.54 (0.14) | **2.06** (0.18) | **0.77** (0.07) | 1.11 (0.10) |
| $\phi^2 = 2$ | SNBM | 21.84 (1.48) | 7.45 (0.72) | 17.90 (0.90) | 4.49 (0.43) |
|  | JNBM | **4.37** (0.33) | **2.15** (0.19) | **3.50** (0.20) | **1.41** (0.11) |
| $\phi^2 = 10$ | SNBM | 176.89 (9.41) | 44.03 (4.33) | 185.50 (6.97) | 26.21 (2.34) |
|  | JNBM | **14.79** (0.69) | **2.20** (0.27) | **16.25** (0.67) | **1.44** (0.13) |

### 3.2 Case Study: the UMICRO Data

We apply JNBM to the UMICRO data set and compare the resulting inferences to those from SNBM in the estimation of regression coefficients and in the prediction of taxa abundance at one of the sites. Again, data were collected from two different body sites by catheterization for urine samples and vaginal swabbing for vaginal samples. Full-length contigs received from Loop Genomics (Element Biosciences, San Diego, CA), which we refer to as LoopSeq, were processed with DADA2 (v 1.24.0) to generate amplicon sequence variants (ASV), using parameters as recommended for synthetic full-length 16S LoopSeq data in [Callahan et al., 2021]. Taxonomic classifications were assigned using BLCA (v 2.2) and the 16S NCBI database (downloaded on 11/16/2021). The processed data contain 68 sample pairs with 151 common taxa (genera) found in both vaginal and urinary samples, while the following analysis focuses on nine taxa of interest: *Gardnerella, Streptococcus, Lactobacillus, Aerococcus, Anaerococcus, Bifidobacterium, Corynebacterium, Fannyhessea, Prevotella*. We incorporate several clinical and demographic metadata for the subjects as covariates in the models: age, body mass index (BMI), diabetes (yes/no), daily yogurt or probiotic consumption (yes/no), race ethnicity (0: White, 1: Black, 2: Others), the presence or absence of overactive bladder (OAB) (yes/no). In addition, the study group variable on menopausal status (Postmenopausal with no estrogen; Postmenopausal on estrogen; Premenopausal) is used to cluster the samples for JNBM SG, introduced in Section 2.2.

Figure 1 shows examples of differences between SNBM and JNBM when it comes to the estimation of regression coefficients. The results indicate that JNBM has relatively pronounced positive effects of BMI and age on *Gardnerella* and *Streptococcus* counts in both vaginal and urinary samples, while these effects are negligible under SNBM. In the previous section, we saw that the joint model provides more accurate estimation results when paired associations are present. Here, the posterior means (and standard deviations) of *ϕ*^2^ for *Gardnerella* and *Streptococcus* are given by 17 (4) and 14 (4), respectively. It favors the results of JNBM for the regression coefficients. Furthermore, the positive effects of BMI and age on the taxa under JNBM are consistent with the biological findings reported in [Brookheart et al., 2019, Xu et al., 2020]. More results on regression coefficients can be found in Supplementary Material A.1.

**Figure 1.**
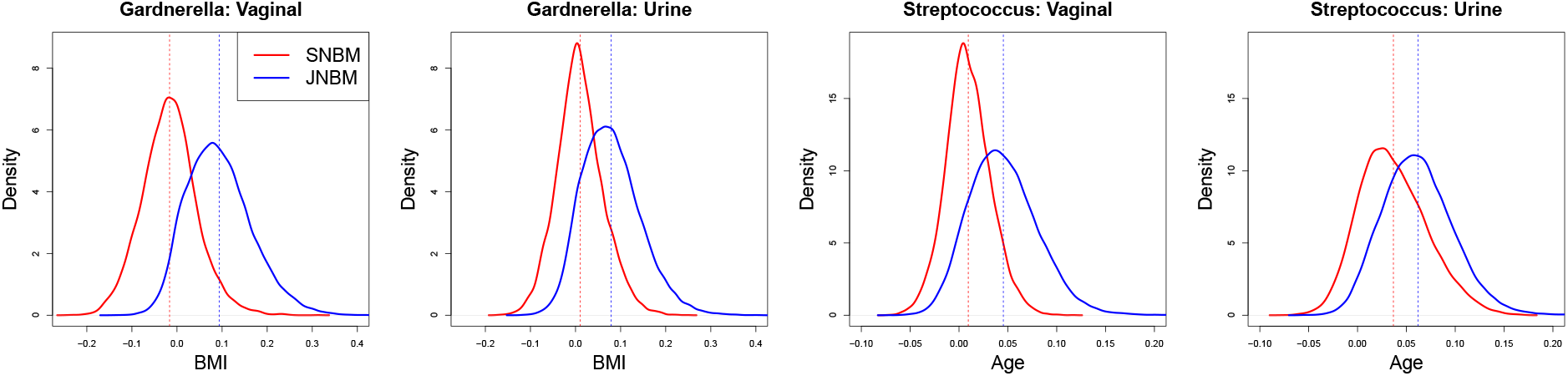
Posterior distributions of regression coefficients for BMI in the vaginal and urine data sets for *Gardnerella* (first two panels); and regression coefficients for Age in the data sets for *Streptococcus* (last two panels). The dashed lines indicate the posterior means.

JNBM SG clusters the samples according to the study group variable, allowing each group to have a different level of paired association. Figure 2 illustrates that, for the three genera, the two postmenopausal groups have similar association strengths but differ from the premenopausal group, which has a weaker association. Figure S11 displays the posterior distributions of the parameters for the nine taxa; interestingly, *Gardnerella* and *Fannyhessea* exhibit remarkable differences in strength among the three groups, with postmenopausal with no estrogen showing the strongest association, followed by postmenopausal with estrogen.

**Figure 2.**
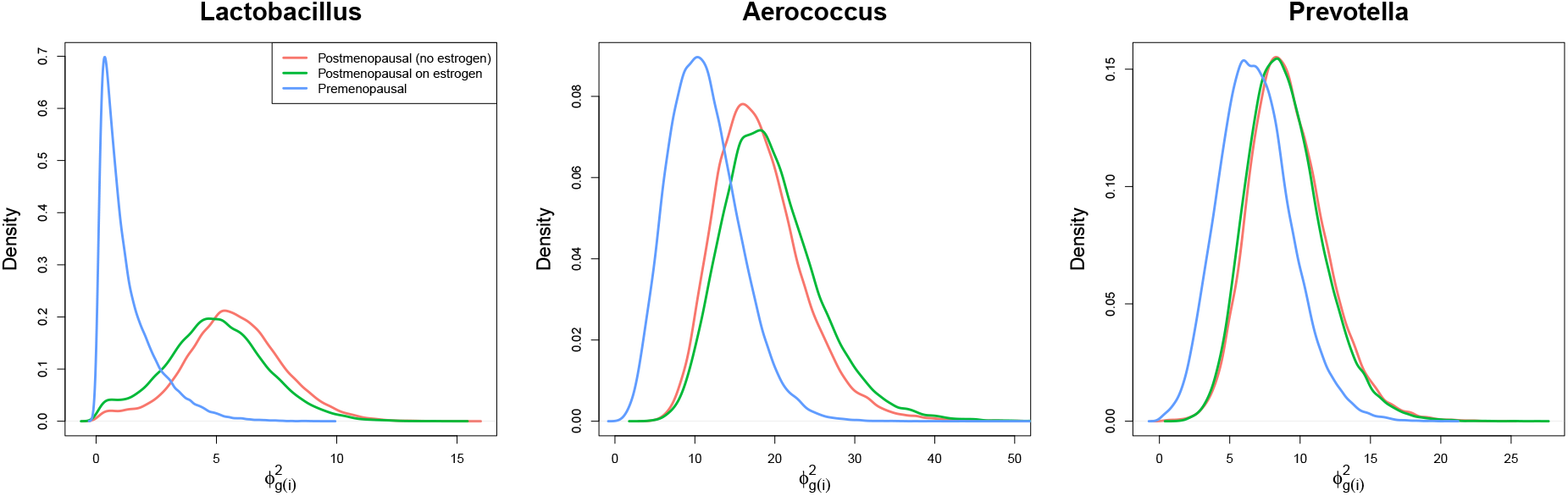
Posterior distributions of 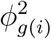, the dispersion hyperparameters of latent factors, from JNBM SG for the three different genera.

To assess the predictive performance of the models for the microbial composition, we calculated the residuals between the observed and predicted relative abundances of each microbial genus, denoted as 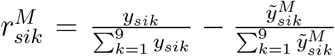, where *y*_*sik*_ is an observed count for the *k*-th taxon in the *i*-th sample of the *s*-th body site and 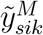 the corresponding predicted count under model *M*. Figure 3 presents the difference in the absolute residuals between SNBM and JNBM SG, that is, 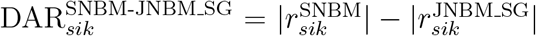. The greater the difference, the better the predictive performance of JNBM SG. The boxplots represent the distributions of DARs per taxon for each body site, with the positive medians (in most taxa) indicating that JNBM SG enhances the predictive efficiency of SNBM. The improvement in prediction under the joint model is attributed to the borrowing of information between sites. As such, the base joint model, JNBM, also outperforms SNBM, which has been further improved by accommodating different levels of association for clusters in JNBM SG (Figure S10) and in JNBM Mix (Figure S12).

**Figure 3.**
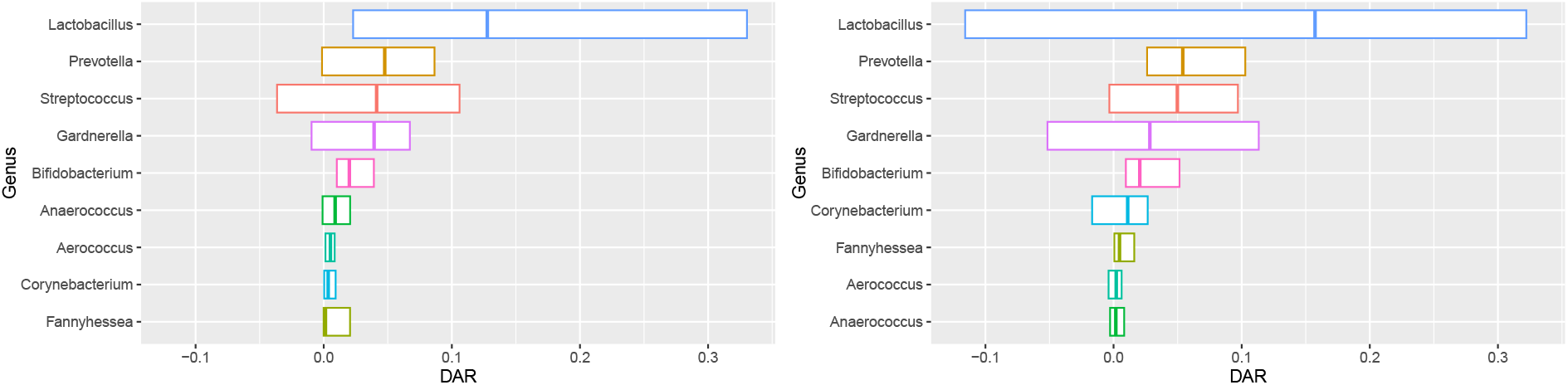
Boxplots (from the 1st to 3rd quartiles) of 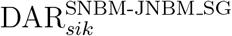, the differences in the absolute residuals between SNBM and JNBM SG, for each taxon. The genera on the y-axis are sorted by their median DARs in each data set.

## 4 Conclusions

We have proposed a joint negative binomial model (JNBM), a latent variable model based on negative binomial distributions, for paired microbiome data. The UMICRO data, gathered from two different body sites for each of the subjects, is a motivating real-world example, and the proposed model incorporates a set of latent factors to capture associations between the two body sites. JNBM has been extended by modifying the distribution for the latent factors, which permits varying levels of paired association across groups of samples.

Our joint negative binomial model provides more accurate regression estimates than the separate negative binomial model without latent factors when there exists a non-zero paired association in the data. In the UMICRO case study, the joint model reveals conspicuous positive effects of BMI and age on *Gardnerella* and *Streptococcus* abundances, respectively, which are in agreement with some previous research findings ([Brookheart et al., 2019, Xu et al.,2020]). Moreover, the joint model outperforms the separate model in prediction, which can be attributed to the use of observations from the opposite body site for prediction via latent factors. With extended models, prediction performance can be further enhanced due to their ability to assign different association strengths to each group.

In addition, estimates of the association strengths can also be used to characterize sample groups. For example, we found that the two postmenopausal cohorts (with and without estrogen) had similar associations but were different from the premenopausal cohort, which had a relatively weaker association, for *Lactobacillus, Aerococcus*, and *Prevotella* in the UMICRO case study.

Although the case study focuses on nine taxa of interest, we have applied these models to other genera. Unsurprisingly, the models struggled to predict taxa that are prevalent in one site but rare in the other. For example, *Escherichia, Klebsiella*, and *Pseudomonas* are pathogens that are common in the urine but scarce in the vagina, showing a high sparsity (high proportion of samples with zero count) in the vaginal data set. It is particularly challenging to predict non-zero counts (urine) using zero counts (vaginal) compared to the reverse scenario. The shared latent factor assumption under our model also becomes questionable when the taxon is rarely present in one of the sites but common in the other.

## Acknowledgement

The research was supported in part by NIGMS grant R01-GM135440, NSF grant EEC-2133504, and NIA grant R03 AG060082.

Supplementary Material for “Negative binomial latent variable model for paired data in microbiome analysis”

## A. Additional figures for real data analysis

### A.1 Posterior distributions of *β*_*s*_

**Figure S1.**
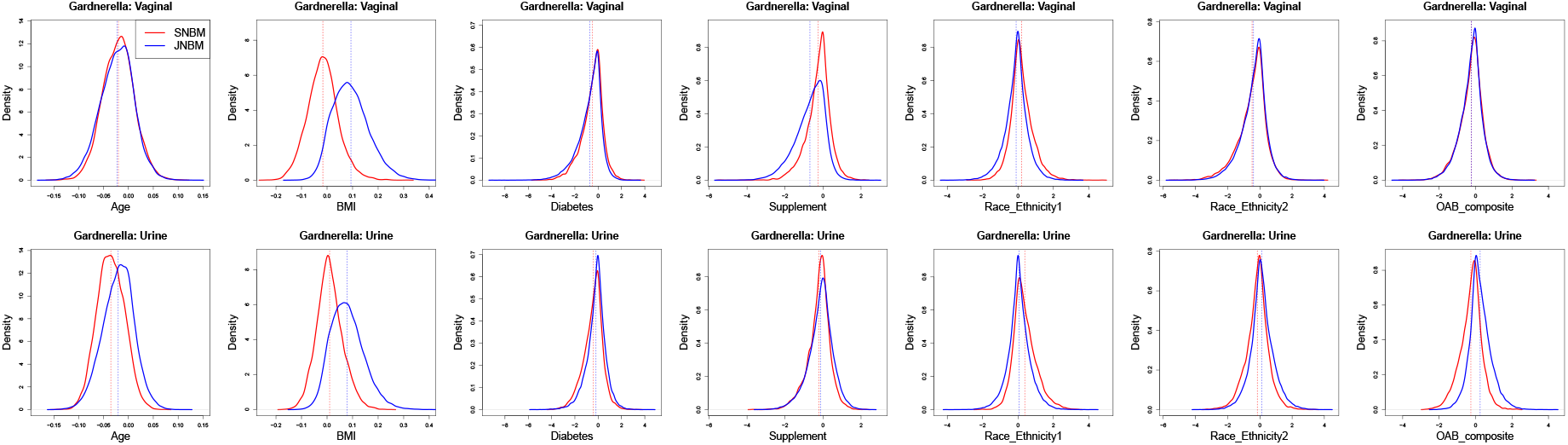
Posterior distributions of regression coefficients in the vaginal (first row) and urine (second row) data sets for *Gardnerella*.

**Figure S2.**
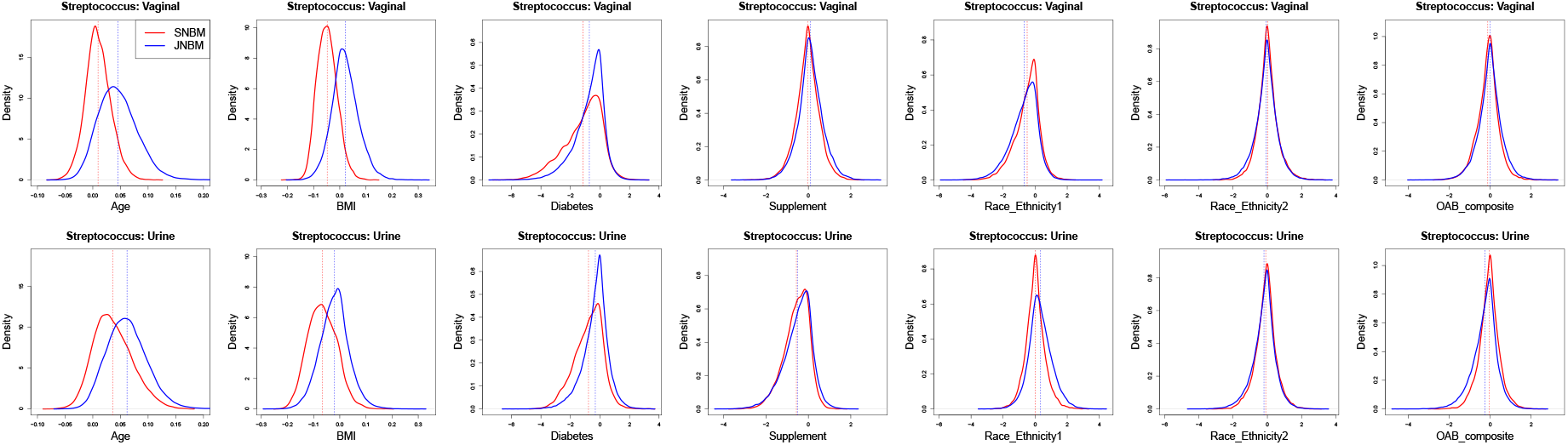
Posterior distributions of regression coefficients in the vaginal (first row) and urine (second row) data sets for *Streptococcus*.

**Figure S3.**
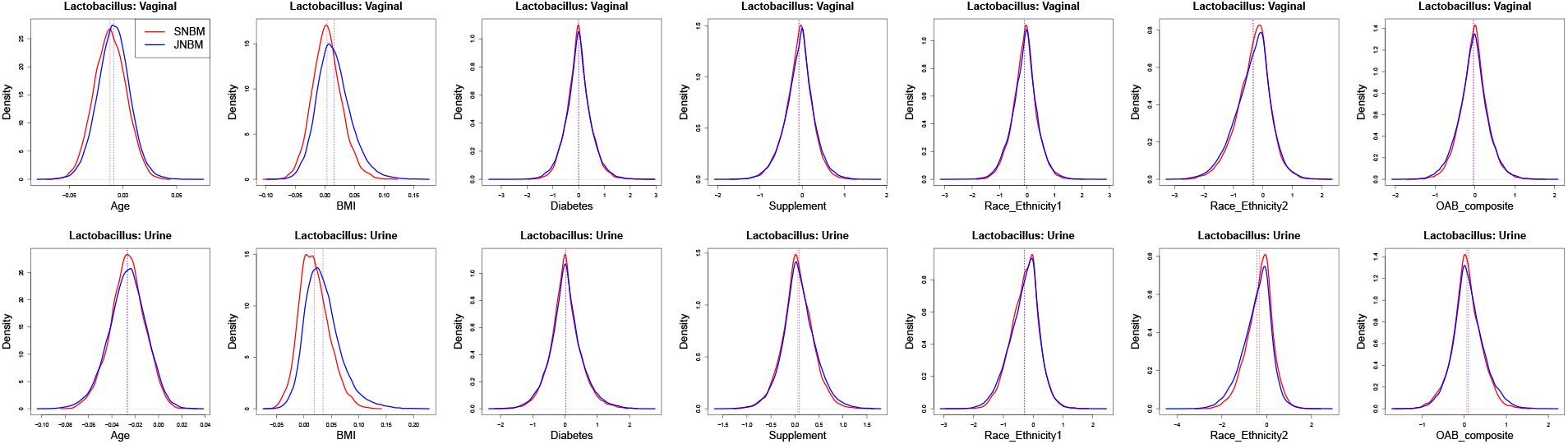
Posterior distributions of regression coefficients in the vaginal (first row) and urine (second row) data sets for *Lactobacillus*.

**Figure S4.**
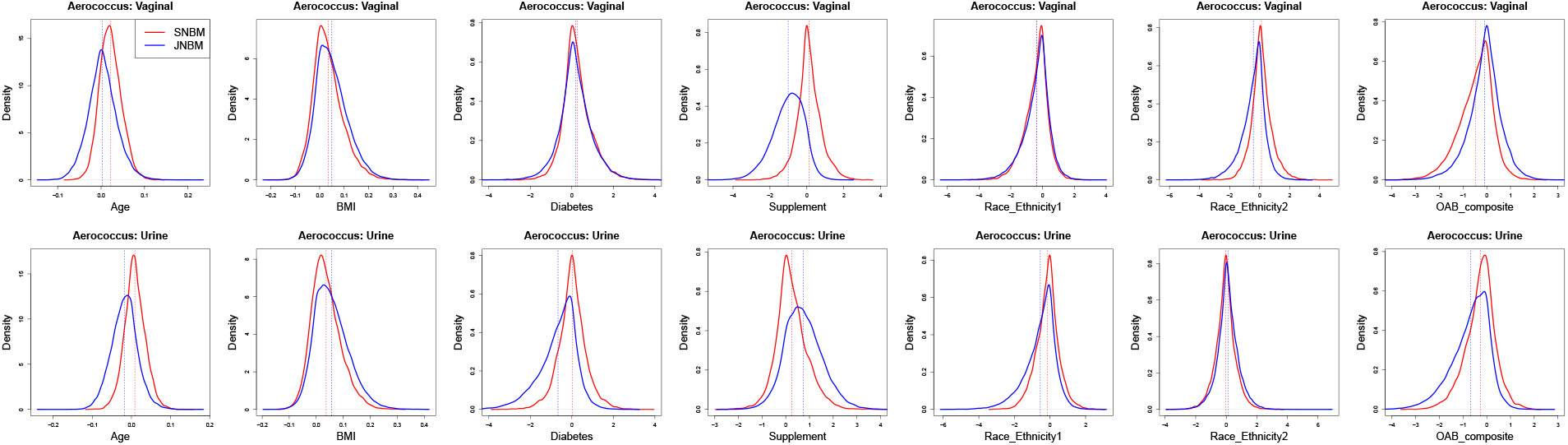
Posterior distributions of regression coefficients in the vaginal (first row) and urine (second row) data sets for *Aerococcus*.

**Figure S5.**
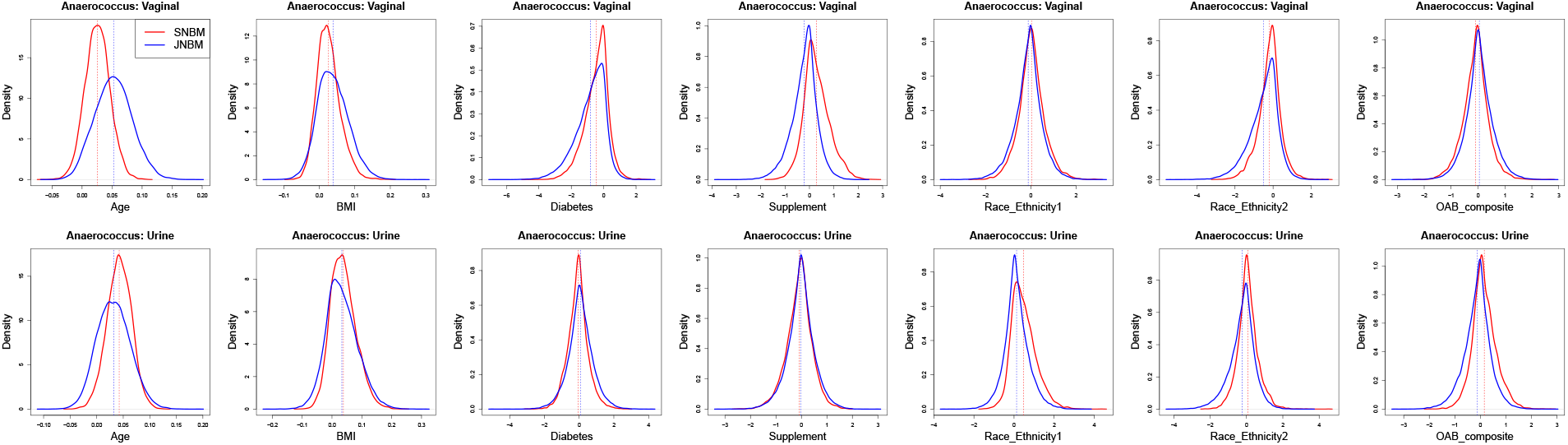
Posterior distributions of regression coefficients in the vaginal (first row) and urine (second row) data sets for *Anaerococcus*.

**Figure S6.**
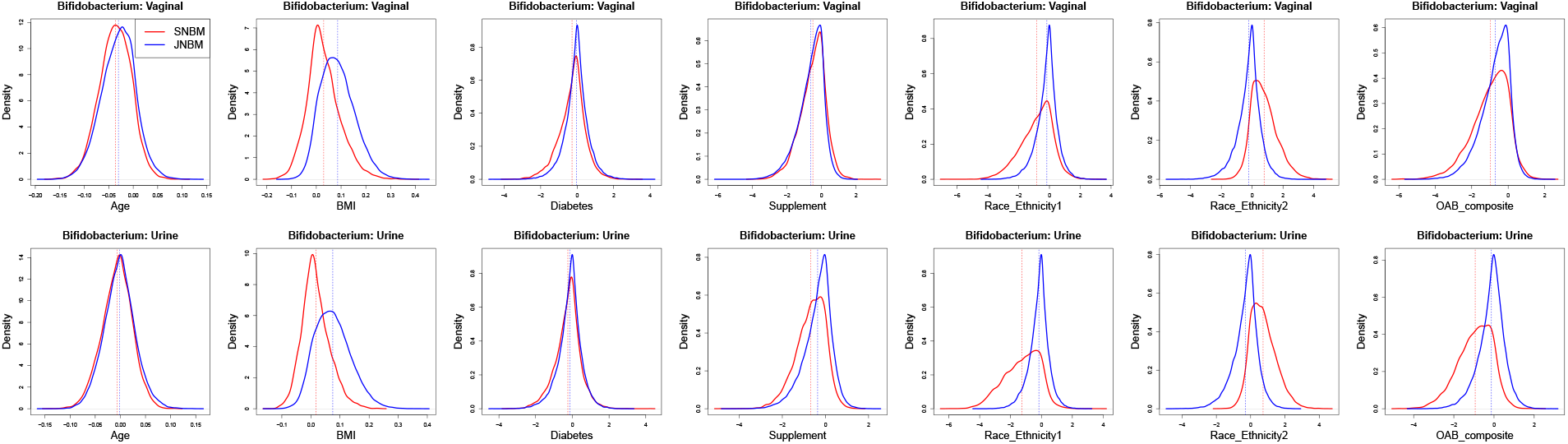
Posterior distributions of regression coefficients in the vaginal (first row) and urine (second row) data sets for *Bifidobacterium*.

**Figure S7.**
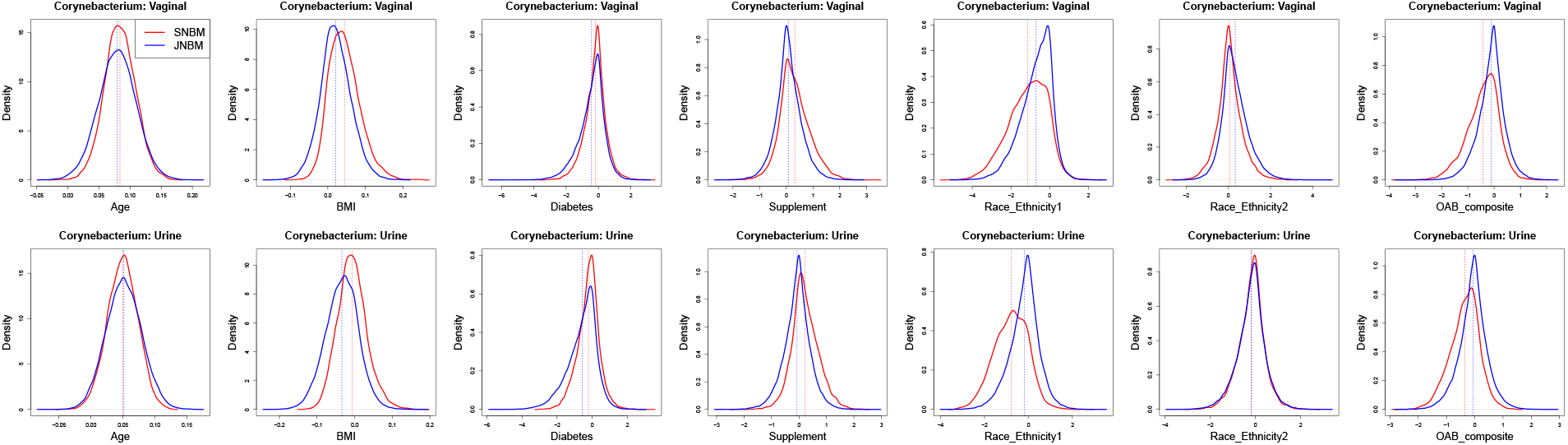
Posterior distributions of regression coefficients in the vaginal (first row) and urine (second row) data sets for *Corynebacterium*.

**Figure S8.**
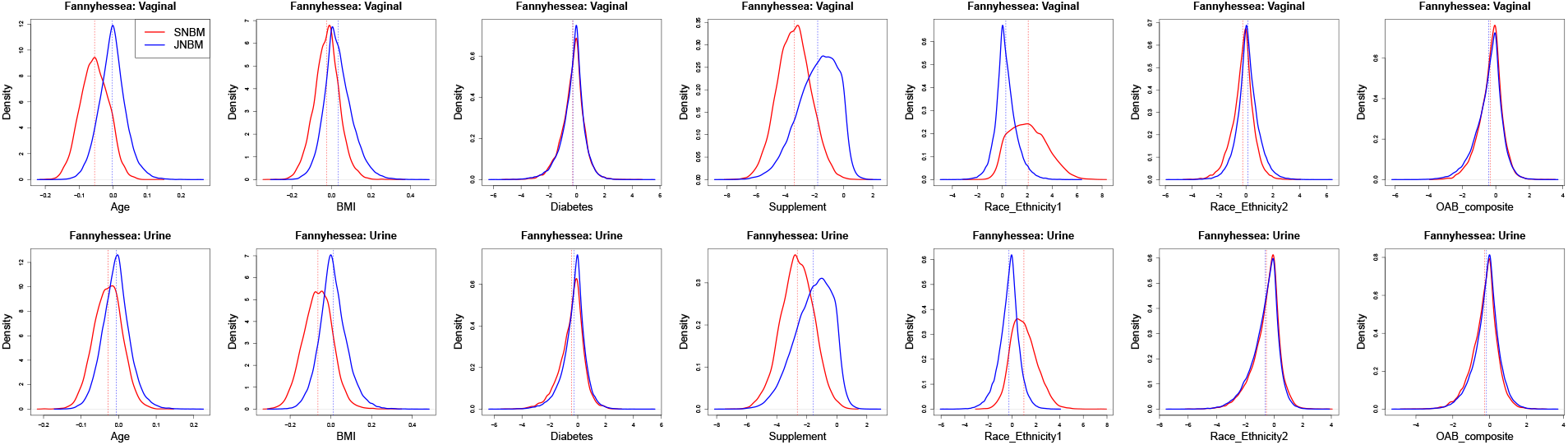
Posterior distributions of regression coefficients in the vaginal (first row) and urine (second row) data sets for *Fannyhessea*.

**Figure S9.**
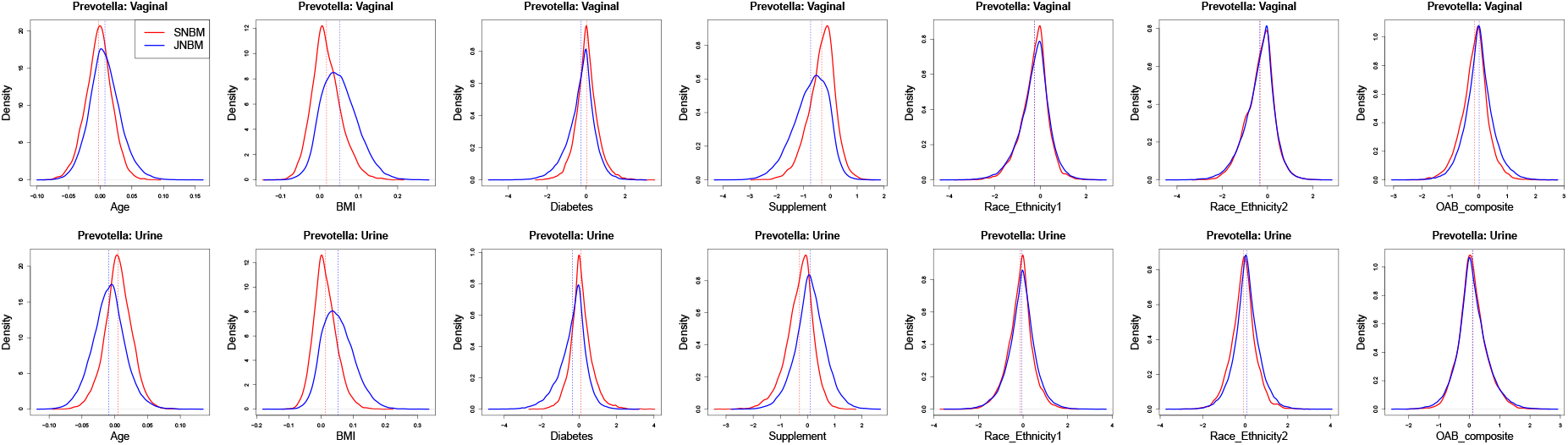
Posterior distributions of regression coefficients in the vaginal (first row) and urine (second row) data sets for *Prevotella*.

### A.2 Boxplots of DARs

**Figure S10.**
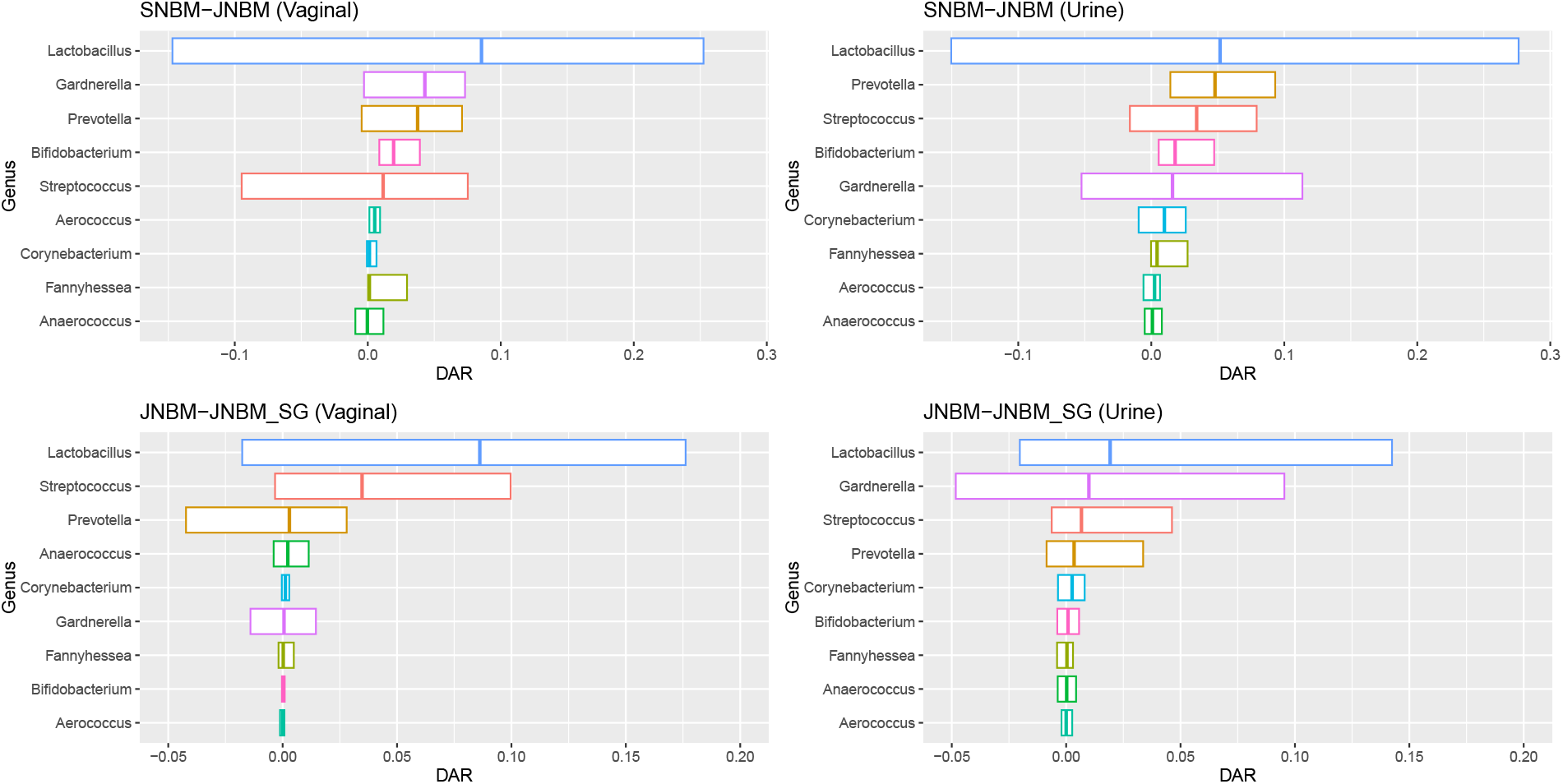
Boxplots of 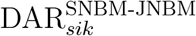 (top row) and 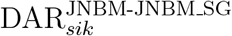 (bottom row).

### A.3 Posterior distributions of 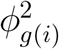 under JNBM SG

**Figure S11.**
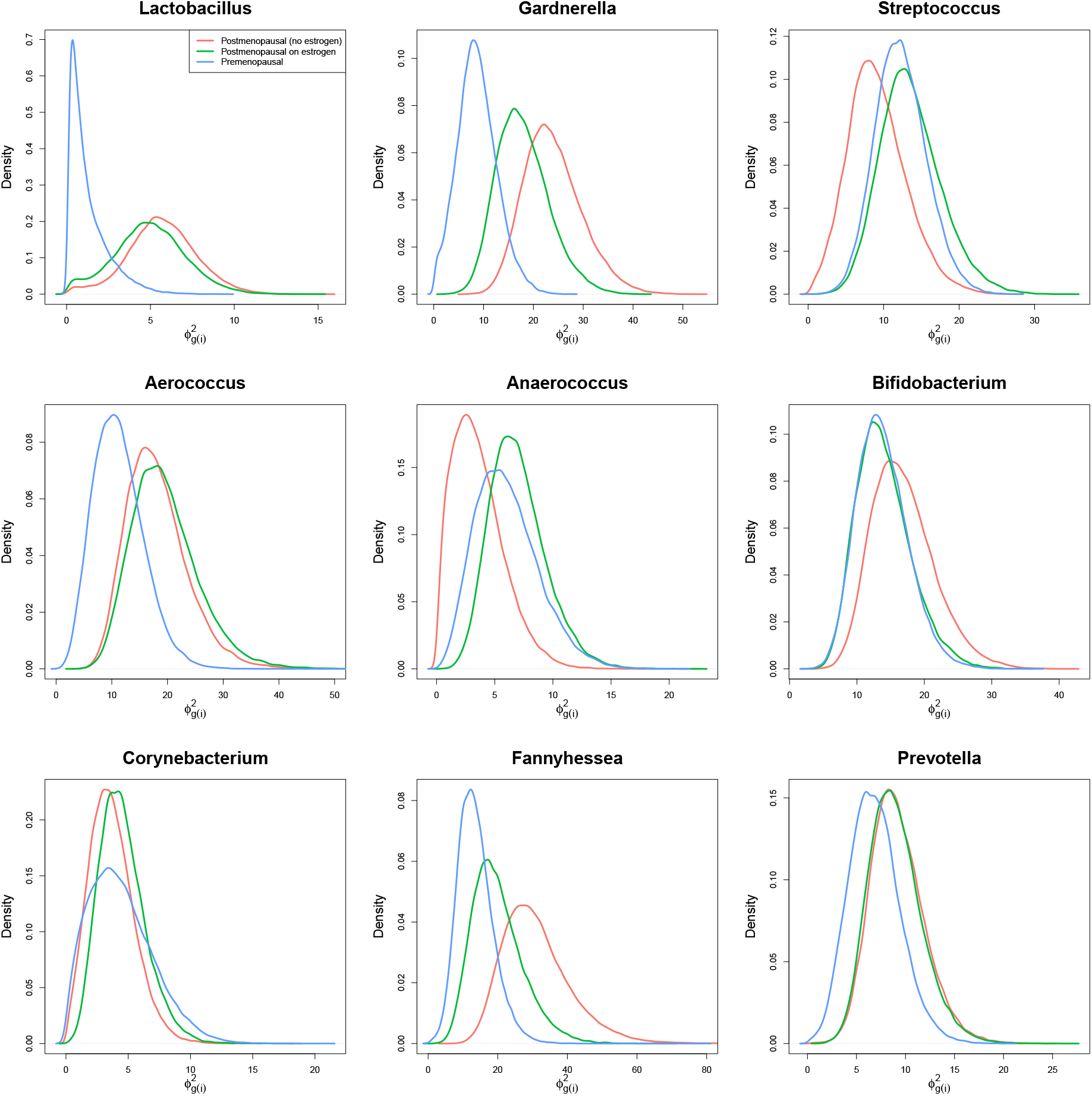
Posterior distributions of 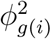 colored by Study Group *g*(*i*).

### A.4 Prediction results of JNBM Mix

**Figure S12.**
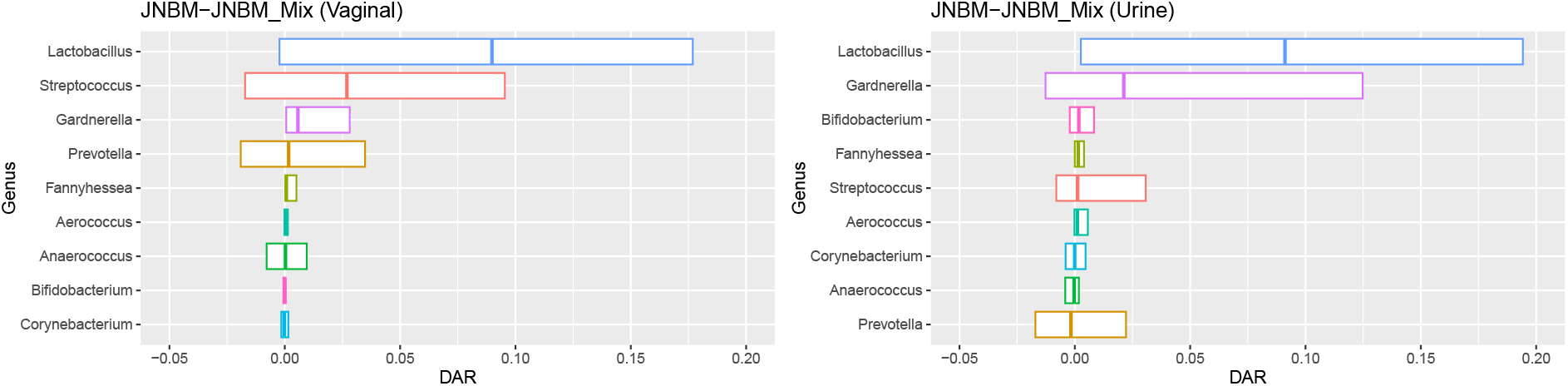
Boxplots of 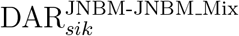.

**Figure S13.**
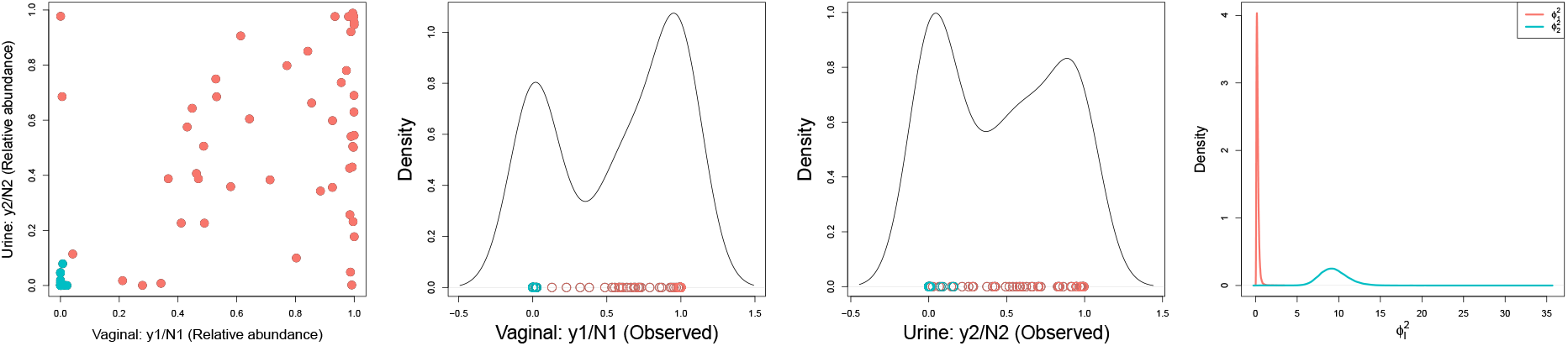
Lactobacillus: On the left is a scatter plot of observed relative abundances colored by posterior estimates of the auxiliary variable *ξ*_*i*_ for the mixture prior; red for 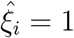 and green for 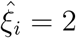, where 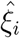 is the posterior median estimate. The next two panels show marginal density estimates of the relative abundances, with observed values at the bottom. The last panel presents the posterior distributions of two dispersion hyperparameters 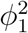 and 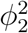.

## B. Computational details on posterior inference

### B.1 Pólya-Gamma augmentation for negative binomial models

This section describes the Pólya Gamma augmentation scheme for the proposed negative binomial model. For sample *i* at body site *s*, the negative binomial probability mass function is given by

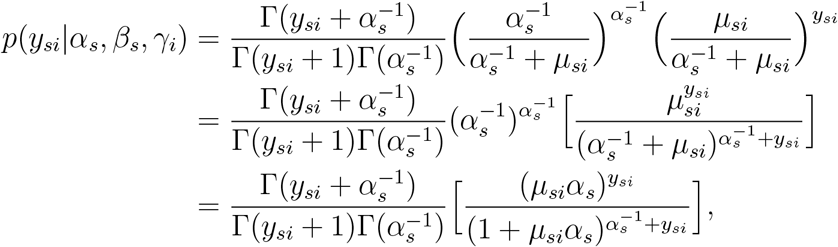

where 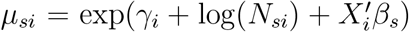 under JNBM. Following [Polson et al., 2013], the last term in the square brackets can be expressed as,

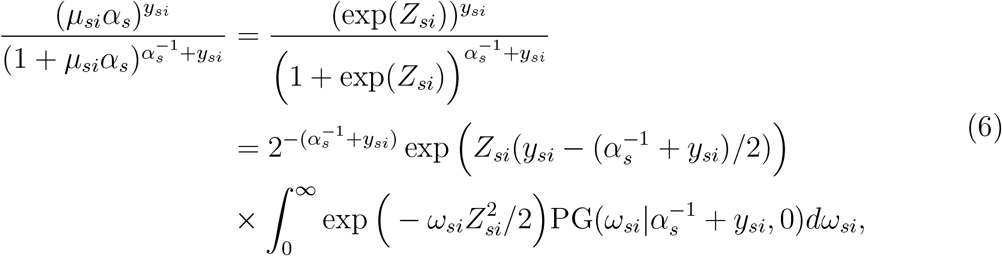

where *Z*_*si*_ *≡* log(*μ*_*si*_) + log(*α*_*s*_). PG(*ω*|*a, b*) denotes the probability density function of the Pólya-Gamma distribution PG(*a, b*). Using the (Pólya-Gamma) auxiliary variable *ω*_*si*_, the equation (6) can be represented hierarchically without the integration. Then, the joint probability function for *y*_*si*_ and *ω*_*si*_ is derived as,

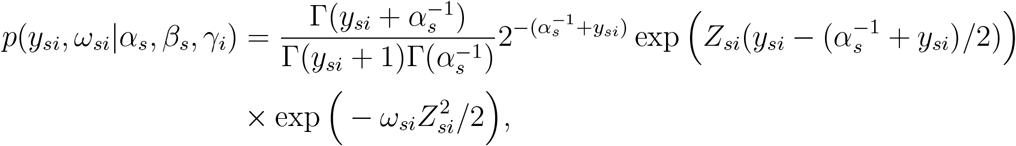

with 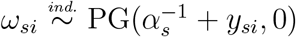 for *i* = 1, ⋯, *n*. The augmented likelihood with the Pólya-Gamma auxiliary variables ensures (normal) prior conjugacy for (*β*_*s*_, *γ*_*i*_), facilitating posterior inference, which will be discussed in the following section.

### B.2 Posterior simulation

The Gibbs sampler is used to draw posterior samples of model parameters for inference. The Pólya-Gamma augmentation enables us to obtain the full conditionals in closed form for most of JNBM parameters except ***α*** = *{α*_1_, *α*_2_*}, ϕ*^2^, and 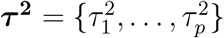, for which we employ the Metropolis–Hastings algorithm. The following are the full conditionals for the parameters of our joint models.

Let ***y***_***s·***_ = *{y*_*si*_ : *i* = 1, ⋯, *n}*, ***γ*** = *{γ*_*i*_ : *i* = 1, ⋯, *n}*, and ***ω***_***s·***_ = *{ω*_*si*_ : *i* = 1, ⋯, *n}* for *s* = 1, 2. With the normal prior 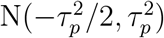, the full conditional for regression coefficients, *β*_*s*_ = (*β*_*s*1_, *β*_*s*2_, ⋯, *β*_*sP*_)^*′*^, is derived as,

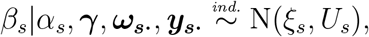

where 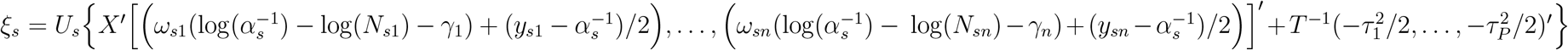 and *U*_*s*_ = (*X*^*′*^Ω_*s*_*X* + *T*^*−*1^)^*−*1^. *X* is a design matrix consisting of the intercept and the covariates for all samples, that is, *X* = (*X*_1_, ⋯, *X*_*n*_)^*′*^ with *X*_*i*_ = (1, *x*_*i*2_, ⋯, *x*_*ip*_)^*′*^. Ω_*s*_ indicates a diagonal matrix of *ω*_*si*_, such that Ω_*s*_ = diag(*ω*_*s*1_, ⋯, *ω*_*sn*_). *T* is also a diagonal matrix of diag 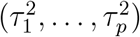.

Similarly, the full conditional for the latent factors ***γ***, with the normal distribution assumption of 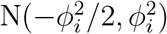 with 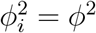, is given by

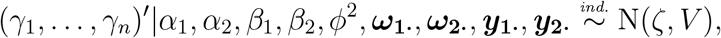

where *V* = (Ω_1_ + Ω_2_ + Φ^−1^)^*−*1^ and 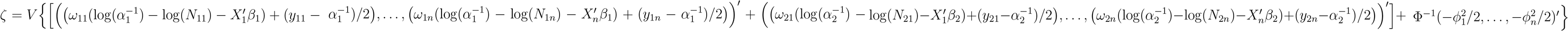. Φ is a diagonal matrix of diag 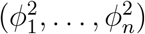.

The full conditional for the auxiliary variables of *ω*_*si*_, with the Polya-Gamma distribution PG 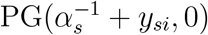 prior, takes the form of

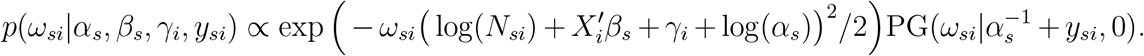

According to Theorem 1 of [Polson et al., 2013],this is proportional to a density function of a Polya-Gamma distribution PG 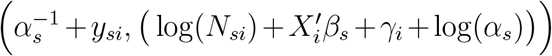; using the prior conjugacy, the posterior samples of *ω*_*si*_ can be taken from the updated Pólya-Gamma distribution.

Other parameters, (***α***, *ϕ*^2^, ***τ*** ^***2***^), have no closed-form full conditionals with the joint full conditional density

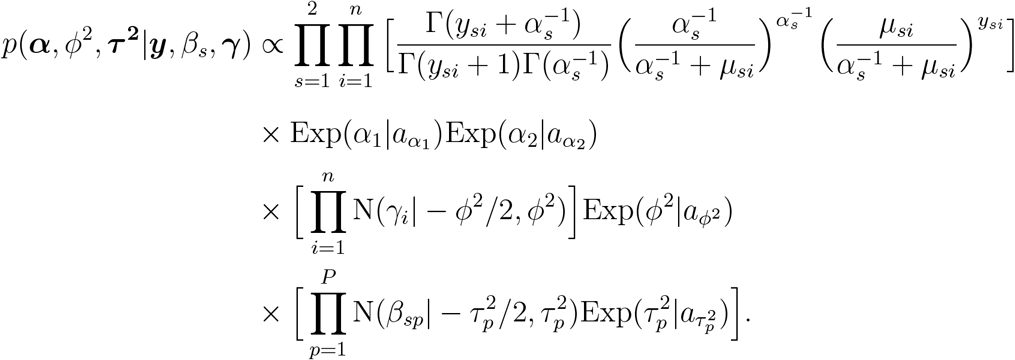

Hence, the parameters are updated with Metropolis-Hastings steps in MCMC, using log-normal proposal distributions.

For the extended models, the dispersion parameters 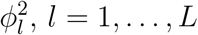, of latent factors are updated using the Metropolis-Hastings algorithm, too. Unlike JNBM SG, JNBM Mix has additional parameters *{ν*_*l*_*}* and *{ξ*_*i*_*}*, given by (4). As Dir 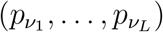 is a conjugate prior for (*ν*_1_, ⋯, *ν*_*L*_), the mixture weight parameters are updated using the Dirichlet distribution with parameters 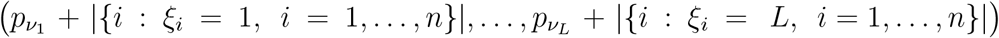, where |*A*| indicates the cardinality of set *A*. Finally, posterior samples of auxiliary variables *{ξ*_*i*_*}* can be drawn from the updated discrete probability function: 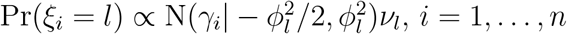, *n* and *l* = 1, ⋯, *L*.

As our modeling strategy is taxon-by-taxon, posterior estimates of model parameters for the microbial community (of multiple taxa) can be obtained by repeating the above Gibbs samplers multiple times, but can be done in parallel.

